# Post-translational knockdown and post-secretional modification of EsxA unambiguously determine the role of EsxA membrane permeabilizing activity in mycobacterial virulence

**DOI:** 10.1101/2020.06.25.170696

**Authors:** Yanqing Bao, Lin Wang, Jianjun Sun

## Abstract

Current genetic studies (e.g. gene knockout) have suggested that EsxA and EsxB function as secreted virulence factors that are essential for *Mycobaterium tuberculosis* (Mtb) virulence, specifically in mediating phagosome rupture and translocation of Mtb to the cytosol of host cells, which further facilitates Mtb intracellular replicating and cell-to-cell spreading. The EsxA-mediated virulence is presumably achieved by its pH-dependent membrane-permeabilizing activity (MPA). However, the data from recent studies have generated a discrepancy regarding to the role of EsxA MPA in mycobacterial virulence with a major concern that genetic manipulations, such as deletion of *esxB-esxA* operon, may stimulate genetic compensation to produce artifacts and/or affect other co-dependently secreted factors that could be directly involved cytosolic translocation. To avoid the drawbacks of gene knockout, we first engineered a *Mycobacterium marinum* (Mm) strain, in which a DAS4+ tag was fused to the C-terminus of EsxB to allow inducible knockdown of EsxB (also EsxA) at the post-translational level. We also engineered a Mm strain by fusing a SpyTag to the C-terminus of EsxA, which allows inhibition of EsxA-ST MPA at the post-secretional level through a covalent linkage to SpyCatcher-GFP. Both post-translational knockdown and post-secretional inhibition of EsxA resulted in attenuation of Mm intracellular survival and virulence in macrophages and lung epithelial cells, which unambiguously confirms the role of EsxA MPA in mycobacterial virulence.

**Author Summary:** Genetic studies, such as loss of function by gene deletion and disruption, have suggested that EsxA is a virulence factor essential for mycobacterial virulence. However, its role is questioned because knockout of *esxA* gene may affect the function or secretion of other related genes. Here, we employed two methods other than gene deletion and disruption to determine EsxA role in mycobacterial virulence. First, we added a degradation signal peptide DAS4+ tag to the C-terminus of EsxB, the chaperon of EsxA so that EsxB-DAS4+ could be degraded by protease ClpXP, whose function can be induced by an inducer, ATC. By this way, we were able to control the amount of EsxB and EsxA at the post-translational level. The results showed that ATC inhibited mycobacterial intracellular survival through down-regulating EsxA and EsxB. Second method is to take advantage of SpyTag(ST) and SpyCatcher(SC) system. Like DAS4+, ST was fused to C-terminus of EsxA without affecting its expression, secretion and MPA. After secretion, EsxA-ST can be specifically recognized by SC-GFP and form a covalent bond between ST and SC, which blocks the MPA, an activity that directly related to mycobacterial virulence. Endogenous expression of SC-GFP in the infected cells inhibited mycobacterial intracellular survival. In summary, our results demonstrate that knockdown of EsxA at the post-translational level or inhibition of EsxA MPA at the post-secretional level, attenuate mycobacterial virulence, and this attenuation is solely attributed to EsxA, not to other factors.

## Introduction

Pathogenic mycobacteria, like tuberculosis and leprosy species, have been imposing great threats to public health for decades (1, 2). Extensive research on their virulence and pathogenicity is urgently needed to minimize the impacts of pathogenic mycobacteria. Completion of whole genome sequencing of multiple mycobacteria species has facilitated researches on mycobacterial pathogenicity, epidemiology, detection and vaccine development (3–9). Various gene editing methods allow researchers to explore genes of interest for their roles in mycobacterial virulence and pathogenicity (10).

The *esxB-esxA* operon is located within the *esx-1* locus in Mtb genome that encodes a Type VII secretion system (11). Current genetic studies (e.g. gene deletion, disruption or mutation) have suggested that EsxA and EsxB play an essential role in mycobacterial pathogenicity, intracellular translocation and escape from immune responses (12–17). EsxA and EsxB are secreted as a heterodimer through the ESX-1 secretion system (18). Our previous studies have demonstrated that EsxA, but not EsxB, exhibits acidic pH-dependent MPA (19). The MPA is uniquely present in the EsxA proteins from pathogenic Mtb and Mm, but is absent in the highly homologous EsxA from non-pathogenic *Mycobacterium smegmatis* (Ms) (19, 20). This suggests that EsxA MPA is the key factor determining the virulence phenotype of mycobacteria. This notion is further confirmed by our recent study showing that single-residue mutations Q5V and Q5K in EsxA either up or down regulated the MPA and consequently up or down regulated the cytosolic translocation and virulence of Mtb and Mm in cultured macrophages and in zebra fish (21). Most recently, we have found that the Nα-acetylation of EsxA at the residue T2 is required for EsxAB heterodimer separation, a prerequisite for EsxA to permeabilize membranes. The non-acetylated mutations at T2 (e.g. T2A and T2R) inhibited the acidic pH-dependent heterodimer separation and consequently attenuated Mm cytosolic translocation and virulence in macrophages (22). Therefore, current studies have established a strong link between EsxA acidic pH-dependent MPA and mycobacterial cytosolic translocation.

While a substantial body of studies have established the link between EsxA MPA and mycobacterial cytosolic translocation, the paradigm has been challenged by two recent reports. In the report from Conrad et. al. (23), the authors successfully repeated our previous in vitro experiments and confirmed that the detergent-free EsxA disrupted liposomal membranes at acidic conditions (19, 20, 24), but they found that Mm was still able to penetrate the phagosome and translocate to the cytosol in the presence of Bafilomycin, a reagent that inhibits intracellular acidification, indicating that phagosome rupture doesn’t occur through the acidic pH-dependent MPA of EsxA (23). Most recently, Lienard et. al. employed a collection of Mm ESX-1 transposon mutants, including the mutants that disrupt EsxA secretion, to infect macrophages and showed that the transposon mutants without EsxA secretion was still able to permeabilize phagosomes, suggesting that other factors independent of EsxAB play a role in cytosolic translocation (17). Therefore, conclusive evidence is required to resolve the discrepancy. It is possible that knockout of the *esxB-esxA* operon stimulates mycobacterial genetic compensatory mechanisms (e.g. secondary mutations and altered gene expression) that produce artifacts (25, 26), and knockout of *esxB-esxA* also affects the co-dependently secreted factor(s) (e.g. EspA, EspC or EspB) that could also play roles in cytosolic translocation (18, 27–30).

In order to determine the exact role of EsxA in mycobacterial virulence, in the present study we employed two approaches to avoid the potential artifacts caused by gene knockout. We first constructed a Mm recombinant strain, namely Mm(EsxB-DAS4+), in which a degradation signal peptide DAS4+ was fused to the C-terminus of EsxB in the genome allowing post-translational knockdown of EsxB upon anhydrotetracycline (ATC) induction (31, 32). Secondly, we engineered another Mm strain, namely Mm(EsxA-ST) by fusing a SpyTag to the C-terminus of EsxA. SpyTag (ST) doesn’t have any noticeable effect on expression, secretion and MPA of EsxA. The ST can be specifically recognized by SpyCatcher (SC), and a covalent amide linkage is automatically formed between a lysine on ST and an asparagine on SC (33, 34). The amide bond formation is fast, irreversible and highly tolerant to various conditions (35). The covalent bonding of SC to ST inhibits the MPA of EsxA-ST at the post-secretional level. The results obtained in the present study have shown that both post-translational knockdown and post-secretional inhibition of EsxA attenuated Mm intracellular survival and virulence.

## Result

### Mm(EsxB-DAS4+) has a reduced secretion and virulence, but it is still significantly more virulent than Mm(ΔEsxA:B)

To avoid the potential artifacts caused by gene knockout, we set off to construct a Mm strain that allows inducible knockdown of EsxA at the post-translational level. Initially, we attached the degradation peptide DAS4+ to the C-terminus of EsxA in the Mm genome. However, EsxA-DAS4+ was not stably expressed in the engineered strain. Since an earlier study has shown that the C-terminal modifications have no significant impact on EsxB’s interaction with other proteins (36), we attached DAS4+ to the C-terminus of EsxB (**Fig. S1A**) and confirmed the construction by PCR (**Fig. S1B**). Western blot analysis showed that EsxB-DAS4+ was expressed in a similar level as EsxB in Mm(WT) in the total lysates. In the culture filtrate, however, EsxB-DAS4+ was less than EsxB, suggesting that the secretion of EsxB-DAS4+ was partly affected by DAS4+ tag (**Fig. 1A**). As expected, the intracellular survival of Mm(EsxB-DAS4+) was ~10 folds lower than that of Mm(WT), but it was still ~100 folds higher than that of Mm(ΔEsxA:B) (**Fig. 1B**).

**Fig. 1.**
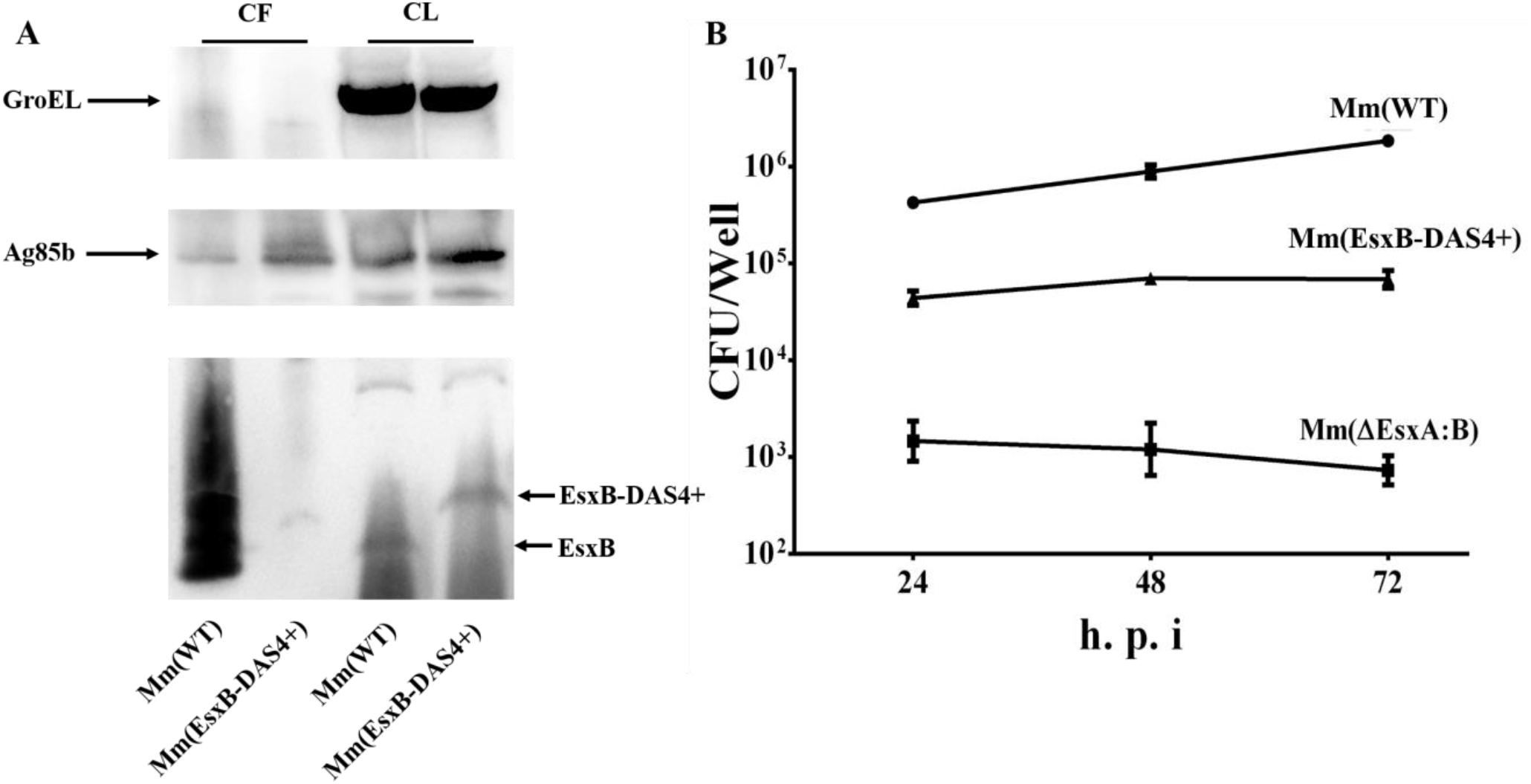
Fusing DAS4+ to the C-terminus of EsxB partly affected EsxB secretion and Mm virulence. (**A**) The culture filtrates (CF) and cell lysates (CL) of Mm(WT) and Mm(EsxB-DAS4+) were applied to SDS-PAGE, and the expression and secretion of EsxB were detected by Western blots by using anti-EsxB serum. As the controls for CF and CL, GroEL and Ag85b were also detected by Western blots using the antibodies against GroEL and Ag85b. (**B**) RAW264.7 cells were infected with Mm(WT), Mm(EsxB-DAS4+) and Mm(ΔEsxA:B) at MOI =1, respectively. At 24, 48 and 72 h of post-infection, the cells were collected and subjected to intracellular survival tests. The CFU of each well was counted to quantify the intracellular survival. The experiment was duplicated and data is presented as mean ± SD.

### Addition of ATC induced knockdown of EsxB-DAS4+ at the post-translational level

To test the inducible knockdown of EsxB-DAS4+, the Mm(EsxB-DAS4+) liquid culture was treated with ATC (0.5 μg/ml) for various times. The lysate was applied to SDS-PAGE, followed by Western blot. The expression of EsxB-DAS4+ was significantly reduced after 6 h of ATC treatment, while ATC had no effect on EsxB even after 48 h treatment on Mm(WT) (**Fig. 2A**). Interestingly, the expression of EsxA was similarly diminished after 6 h treatment (**Fig. 2A**). Given that EsxA and EsxB reportedly form a heterodimer, the protease may induce degradation of the heterodimer at the post-translational level. Moreover, ATC did not down-regulate the transcription level of *esxB-DAS4+* (**Fig. 2B**), indicating the inducible knockdown specifically targets to protein. Considering the limited sensitivity of the antibodies used in Western blot, immunofluorescence assay was used to further determine the ATC-induced knockdown of EsxB-DAS4+ in Mm(EsxB-DAS4+). As expected, at 48 h of ATC treatment, EsxB-DAS4+ was significantly reduced (**Fig 2C, D**). Interestingly, EsxA was also significantly reduced (**Fig 2E, F**), which is consistent to the finding that EsxA and EsxB form a heterodimer.

**Fig. 2.**
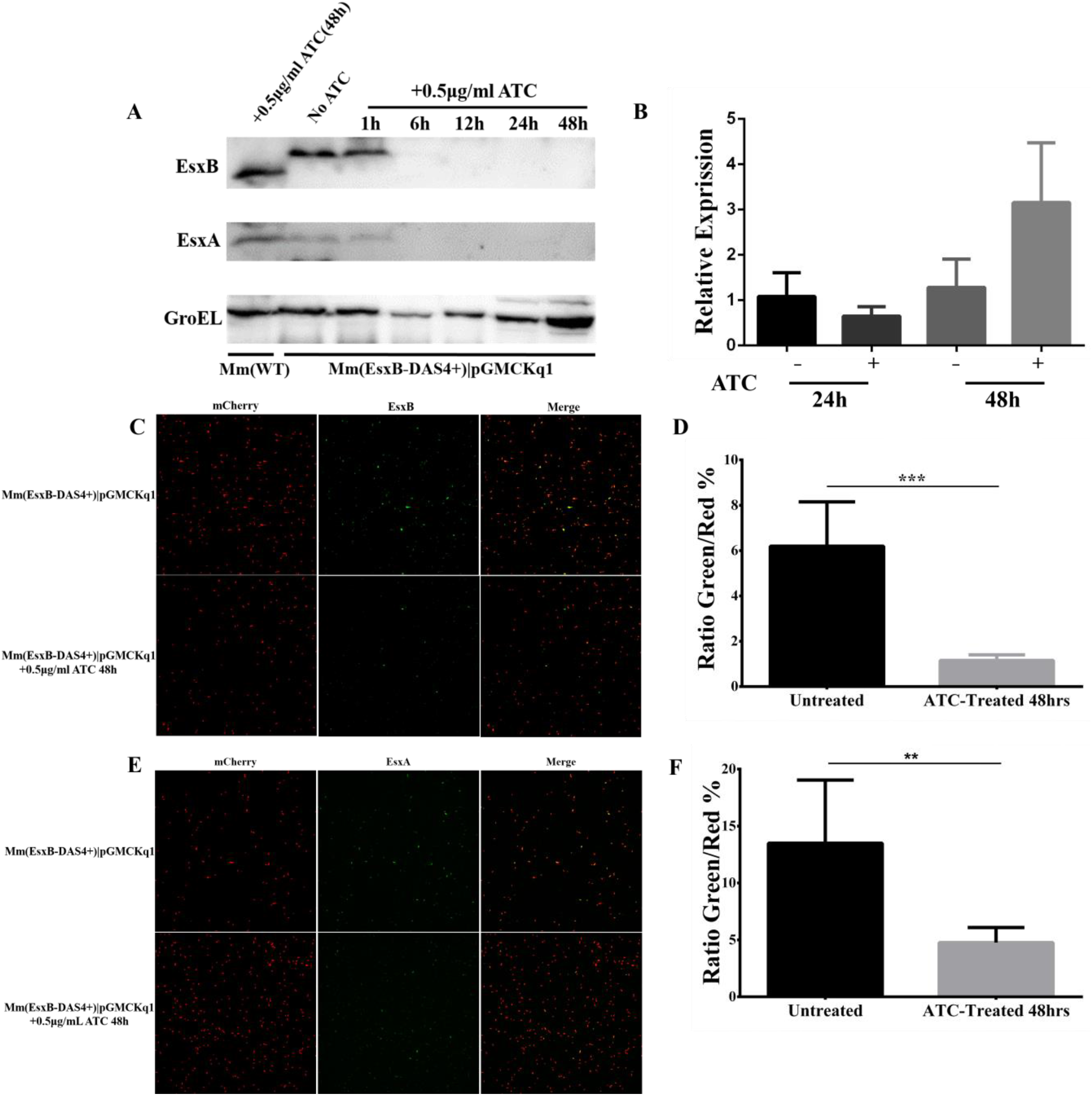
ATC induced knockdown of EsxB-DAS4+ at the post-translational level. (**A**) The Mm(EsxB-DAS4+)|pGMCKq1 culture was treated without ATC or with ATC (0.5 μg/ml) for the indicated times. As a control, the Mm(WT) culture was also treated with ATC (0.5 μg/ml) for 48 h). The expression of EsxB-DAS4+, EsxA and GroEL were detected by Western blots using the antibodies against EsxB, EsxA and GroEL, respectively. (**B**) The relative level of EsxB-DAS4+ mRNA after 24 h and 48 h of ATC treatment was determined by RT-qPCR. The data was presented as mean ± SD and analyzed with multiple *t* tests. (**C**) The mCherry-expressing Mm(EsxB-DAS4+)|pGMCKq1 cells were treated without or with ATC (0.5 μg/ml) for 48 h. Then the bacteria were incubated with anti-EsxB serum, followed by FITC-labeled secondary antibody, to label the surface-associated EsxB-DAS4+. The images were taken under a LSM700 confocal fluorescence microscopy. (**D**) The Green/Red ratio in the randomly selected fields was quantified. (**E**) The mCherry-expressing Mm(EsxB-DAS4+)|pGMCKq1 cells were treated without or with ATC (0.5 μg/ml) for 48 h. Then the bacteria were incubated with anti-EsxA serum, followed by FITC-labeled secondary antibody, to label the surface-associated EsxA. The images were taken under a LSM700 confocal fluorescence microscopy. (**D**) The Green/Red ratio in 5 randomly selected fields was quantified. The IFA experiments were duplicated and data is present as mean ± SD. The statistical analysis was performed with *t* test. \*\**P*<0.01, \*\*\**P*<0.001.

### ATC-induced knockdown of EsxB-DAS4+ attenuated Mm’s intracellular survival in mammalian cells

We then tested the effects of ATC induction on Mm’s intracellular survival in WI-26 cells. The Mm(EsxA-DAS4+)|pGMCKq1 cells were pre-treated with ATC (0.5 μg/mL) before infection and then were applied to infection of WI-26 cells. As expected, the ATC-treated Mm(EsxB-DAS4+)|pGMCKq1 had a significantly lower intracellular survival than that without ATC treatment. As controls, the intracellular survival of Mm(WT) is not affected by ATC (**Fig. 3A**). Next, we set out to test the effects of ATC treatment during the infection on Mm intracellular survival. First, we titrated the cytotoxic effect of ATC and found that ATC did not have significant cytotoxicity at a concentration up to 5 μg/mL (**Fig. 3B**). To ensure knockdown of EsxB when Mm(EsxB-DAS4+) is inside the host cell, the highest concentration without cytotoxicity (5 μg/mL) was applied to cell culture medium at the time of infection (0 hpi). At 24 hpi and 48 hpi, the intracellular survival of Mm(EsxB-DAS4+)|pGMCKq1 with ATC addition was significantly lower than the group without ATC (**Fig. 3C**). ATC had no effect on the intracellular survival of Mm(WT) at 24 hpi, but had a down-regulatory effect at 48 hpi. It could be because that ATC is an antibiotic that inhibits bacterial growth overtime, or it has cytotoxic effects to the host cells. However, we noticed that the ATC unspecific inhibition to Mm(WT) was much less than its specific inhibition for Mm(EsxB-DAS4+) (**Fig. 3C**). Similar results were acquired from THP-1 cell line (**Fig. 3D**). Together, knockdown of EsxB-DAS4+ by ATC either before or during the infection attenuates mycobacterial intracellular survival.

**Fig. 3.**
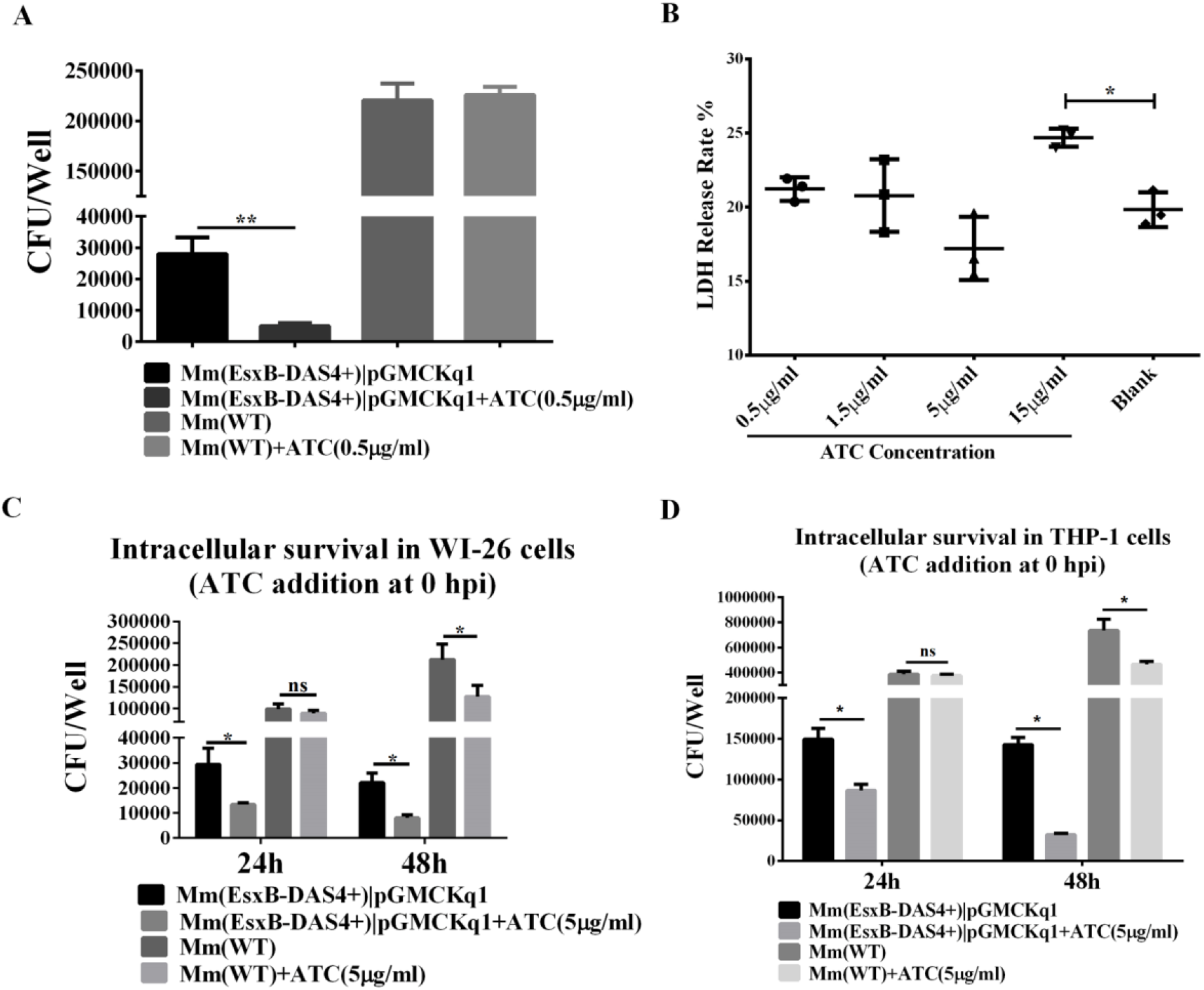
Inducible knockdown of EsxB reduced Mm intracellular survival. (**A**) Mm(EsxB-DAS4+)|pGMCKq1 and Mm(WT) were treated with/out ATC (0.5 μg/ml) for 48 h. Then WI-26 cells were infected with the indicated mycobacteria at MOI=1. At 24 hpi, the cells were collected and subjected to intracellular survival assay (CFU/well). (**B**) WI-26 cells were treated with ATC at various concentrations for 24 h. The cytotoxicity was measured by LDH release assay. (**C**) WI-26 cells were infected with Mm(EsxB-DAS4+)|pGMCKq1 and Mm at MOI=1. At the time of infection (0 hpi), ATC (5 μg/ml) was added to the cell culture to induce EsxB-DAS4+ knockdown. At 24 and 48 hpi, the cells were harvested and subjected to intracellular survival assay. (**D**) Similarly, THP-1 cells were infected with Mm(EsxB-DAS4+)|pGMCKq1 and Mm at MOI=1. At the time of infection (0 hpi), ATC (5 μg/ml) was added to the cell culture to induce EsxB-DAS4+ knockdown. At 24 and 48 hpi, the cells were harvested and subjected to intracellular survival assay. The experiments were duplicated and data is present as mean ± SD. For cytotoxicity data, the statistical analysis was performed with One-way ANOVA, followed by Holm-Sidak multiple comparison. For CFU data, the statistical analysis was performed with multiple *t* test between ATC treated and nontreated groups of each strain. \**P*<0.05, \*\**P*<0.01.

### Insertion of ST to the C-terminus of EsxA allows SC-GFP to covalently modify EsxA-ST at the post-secretional level

According to an early report, modification of EsxA N-terminus impaired its secretion and function (37), so we engineered the ST to C-terminus of EsxA with the suicide plasmid (**Fig. S2**). The insertion of ST was confirmed by PCR using the overlap primers (**Fig. S2**). To test if EsxA-ST is expressed, the lysate of Mm(EsxA-ST) was incubated with the purified SC-GFP and then applied to SDS-PAGE, followed by Western blot using either anti-EsxA antibody (**Fig. 4A**) or anti-GFP antibody (**Fig. 4B**). The results showed that EsxA-ST was successfully expressed, and a portion of EsxA-ST reacted with SC-GFP to form a higher molecular weight complex EsxA-ST-SC-GFP (~ 70 kDa) (**Fig. 4A and B**). As a control, SC-GFP did not react with EsxA in the lysate of Mm(WT). Next, we tested if SC-GFP reacts with the bacterial surface-associated EsxA-ST. The live mCherry-expressing Mm(EsxA-ST) and Mm(WT) were first incubated with SC-GFP, and then the cells were washed to remove free SC-GFP, which was followed fluorescence microscopy. We found that Mm(EsxA-ST) was labeled by SC-GFP, but Mm(WT) was not, indicating that SC-GFP specifically recognized EsxA-ST and formed the EsxA-ST-SC-GPF complex on the cell surface (**Fig. 4C**). Moreover, Mm(EsxA-ST) had a similar intracellular survival as Mm(WT), suggesting that ST doesn’t affect expression, secretion and function of EsxA (**Fig. 4D**).

**Fig. 4.**
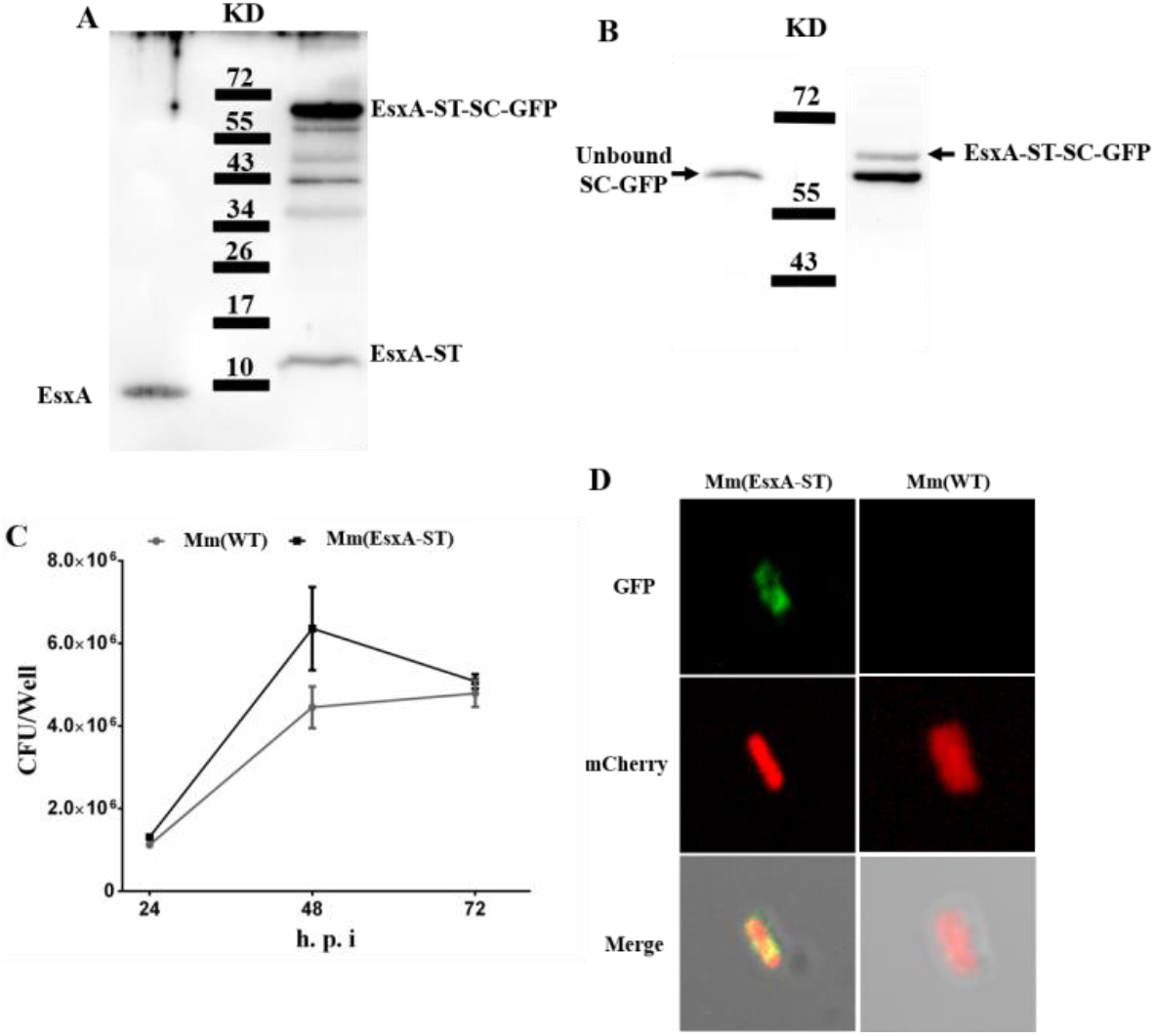
Insertion of SpyTag (ST) at the C-terminus of EsxA doesn’t affect Mm virulence and allows post-secretion labeling of EsxA by SpyCatcher(SC)-GFP. (**A**) The total lysates of Mm and Mm(EsxA-ST) were incubated with the purified SpyCatcher(SC)-GFP and then subjected to SDS-PAGE. EsxA, EsxA-ST and EsxA-ST-SC-GFP were detected by Western blot using anti-EsxA serum. (**B**) Subsequently, the free SC-GFP and EsxA-ST-SC-GFP were detected by Western blots using anti-GFP antibody. (**C**) RAW264.7 cells were infected with Mm or Mm(EsxA-ST) at MOI=1. At 24, 28 and 72 hpi, the cells were collected and subjected to the intracellular survival assay. The experiments were duplicated and data is presented as mean ± SD. (**D**). The mCherry-expressing Mm and Mm(EsxA-ST) were incubated with the purified SC-GFP. After washes, the mycobacterial cells were subjected to fluorescence microscopy. The images were taken at red channel and green channel, respectively.

### SC-GFP inhibited the MPA of EsxA-ST in liposomes

Here, we determined the effect of SC-GFP on MPA of EsxA-ST using the ANTS/DPX dequenching assay in liposome as previously described (19) (**Fig. 5**). At pH 4, EsxA-ST and EsxA induced similar liposome leakage, confirming that ST does not affect the MPA. However, in the presence of SC-GFP, the MPA of EsxA-ST was significantly inhibited. As controls, both EsxA and EsxA-ST were not active in membrane disruption at pH 7.

**Fig. 5.**
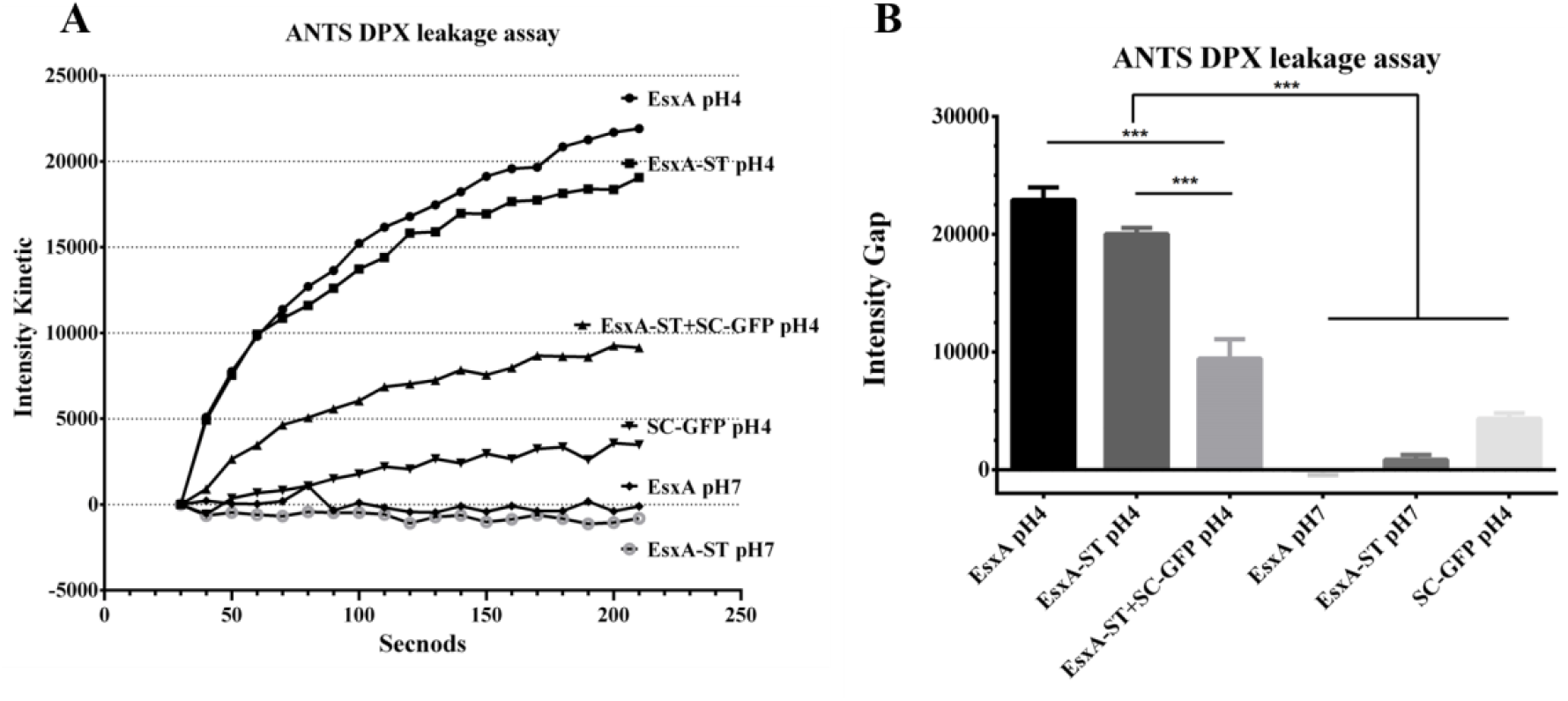
SC-GFP inhibited membrane-permeabilizing activity of EsxA-ST in the liposome leakage assay. (**A**) The liposomes containing ANTS (fluorophore) and DPX (quencher) were incubated with the indicated proteins at either pH 7 or pH 4. The dequenching of ANTS fluorescence was recorded with times. (**B**) The relative fluorescence intensity change between 30 s and 210 s was calculated for each group. The experiment was duplicated and data is presented as mean ± SD. For column graph, statistical analysis was performed with One-way ANOVA. \*\*\**P*<0.001.

### Endogenous expression of SC-GFP inhibited Mm(EsxA-ST) intracellular survival

To determine whether SC-GFP inhibits Mm(EsxA-ST) virulence, we endogenously expressed SC-GFP by transient transfection in human type I lung epithelial cell A549 cells (**Fig. S3**). Cellular fractionation analysis showed that the majority of SC-GFP is localized in the cytosol and a minor portion is localized in the membrane fraction. The A549 cells were transiently transfected with SC-GFP for 24 h and then infected with Mm(EsxA-ST) and Mm(WT), respectively. Since both transfection efficiency and infection efficiency are less than 100% and vary, in order to obtain accurate and reliable results, we measured the intracellular survival with two independent approaches, fluorescence microscopy and CFU counting. In the fluorescence microscopy method, we selected the cells containing both mCherry fluorescence (mycobacterial cells) and green fluorescence (either GFP or SC-GFP) and then quantified the Red/Green ratio in each cell (**Fig. 6A**).

**Fig. 6.**
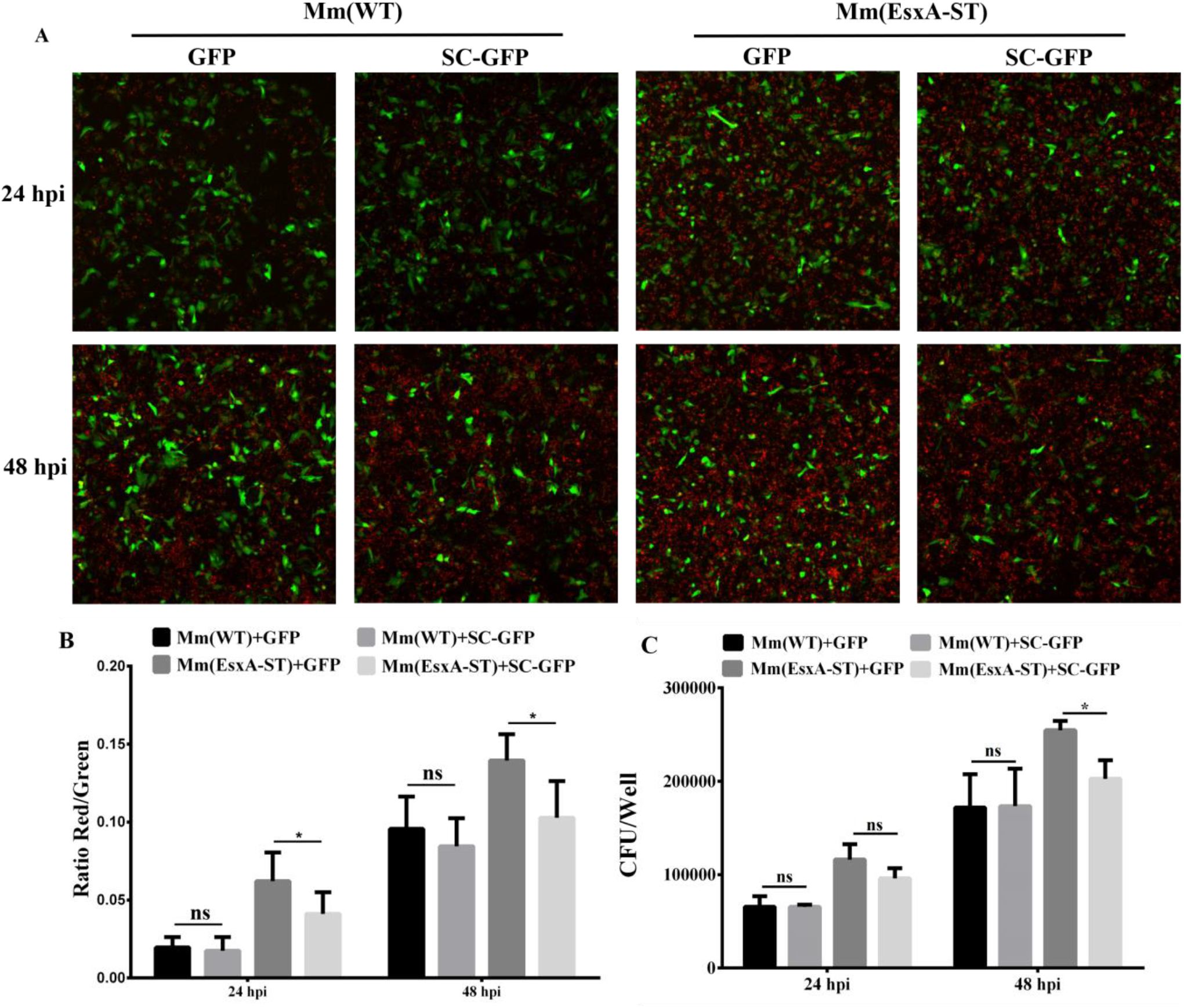
Endogenous expression of SC-GFP reduced Mm(EsxA-ST) intracellular survival in lung epithelial cells. (**A**) A549 cells were transiently transfected with pcDNA3.1-GFP or pcDNA3.1-SC-GFP. After 24 hours of transfection, A549 cells were infected with Mm(WT) and Mm(EsxA-ST) at MOI=2, respectively. At 24 and 48 hpi, the cells were fixed and images were taken at green channel and red channel to visualize GFP/SC-GFP and mCherry-expressing bacteria. (**B**) The Red/Green ratio of the cells in randomly selected fields was calculated to assess bacterial intracellular survival. (**C**) A549 cells were transiently transfected with pcDNA3.1-GFP or pcDNA3.1-SC-GFP. After 24 hours of transfection, A549 cells were infected with Mm and Mm(EsxA-ST) at MOI=2, respectively. At 24 and 48 hpi, the cells were harvested and subjected to CFU counting for intracellular survival. The experiment was duplicated and data is presented as mean ± SD. Statistical analysis was performed with multiple *t*-test between GFP and SC-GFP groups of each strain. \**P*<0.05.

The Red/Green ratio of the cells containing Mm(EsxA-ST) and SC-GFP was significantly lower than that of the cells containing Mm(EsxA-ST) and GFP (**Fig. 6B**), while the Red/Green ratio of the cells containing Mm(WT) and SC-GFP was similar to the cells containing Mm(WT) and GFP (**Fig. 6B**), suggesting that SC-GFP specifically inhibited Mm(EsxA-ST) intracellular survival. Consistent results were obtained in the CFU counting assay (**Fig. 6C**), showing that at 48 hpi, the intracellular survival of Mm(EsxA-ST) in the cells expressing SC-GFP was significantly lower than that in the cells expressing GFP.

## Discussion

Despite of extensive studies in the past decades, the data regarding to the role of EsxA in Mtb pathogenesis, particularly in phagosome rupture and cytosolic translocation, is still conflicting. The artifacts resulted from gene knockout may be a major reason for the discrepancy and confusion. One, knockout of the *esxB-esxA* operon may stimulate hosts to produce compensatory mechanisms, such as secondary mutations or alternations of expression of other genes. Two, knockout of EsxAB will affect other factors (e.g. EspA, EspB and EspC) that are co-dependently secreted with EsxAB. Therefore, to avoid the potential artifacts induced by gene deletion, we engineered the Mm(EsxB-DAS4+) strain to allow inducible knockdown of EsxB (also EsxA) at the post-translational level, which presumably minimizes potential compensatory mutations (**Figs. 1–2**). As expected, inducible knockdown of EsxB significantly attenuated Mm intracellular survival (**Fig. 3**). However, there is still a concern that inducible knockdown of EsxB and EsxA affects the secretion of other co-dependent factors. Therefore, we constructed Mm(EsxA-ST) stain to allow inhibition of EsxA MPA at the post-secretion level. The data showed that attaching ST to the C-terminus of EsxA does not affect the expression, secretion and function of EsxA, and hence it has no effect on Mm virulence (**Fig. 4**). The MPA of EsxA-ST can only be specifically inhibited by SC-GFP through a covalent bonding (**Fig. 5**). Endogenous expression of SC-GFP attenuated the intracellular survival of Mm(EsxA-ST) (**Fig. 6**). Therefore, the data in this study conclusively support that EsxA MPA plays a direct role in mycobacterial intracellular survival and virulence.

In fact, the data obtained in this study is highly consistent with our earlier reports. We have found that single-residue mutations Q5V and Q5K in EsxA either up or down regulated the MPA and consequently up or down regulated the cytosolic translocation and virulence of Mtb and Mm in cultured macrophages and in zebra fish (21). Most recently, we have found that the Nα-acetylation of EsxA at the residue T2 is required for EsxAB heterodimer separation, a prerequisite for EsxA to permeabilize membranes. The non-acetylated mutations at T2 (e.g. T2A and T2R) inhibited the acidic pH-dependent heterodimer separation and consequently attenuated Mm cytosolic translocation and virulence in macrophages (22). Taken together, we conclude that EsxA mediates phagosome rupture and mycobacterial cytosolic translocation through the pH-dependent MPA. The DAS4+ system is based on mycobacterial innate protease ClpXP to degrade the intracellular protein (31, 32). Since ClpXP’s protease expression is dependent on adaptor protein SspB, the inducible SspB expression makes the degradation conditionally (38, 39). Initially, the DAS4+ tag was attached to C-terminus of EsxA, but EsxA-DAS4+ was not stable, so we engineered Mm(EsxB-DAS4+). While EsxB-DAS4+ was stable, it had a lower secretion than Mm (WT) EsxB (**Fig. 1A**). Thus, Mm(EsxB-DAS4+) had a lower virulence than Mm(WT), but it is still much stronger than Mm(ΔEsxA:B) (**Fig. 1B**). Interesting, ATC induction also caused degradation of EsxA, which is consistent to the fact that EsxB and EsxA form a heterodimer (40). Moreover, ATC did not affect the mRNA level of EsxB. Together, ATC-induced knockdown of EsxB (also EsxA) occurs at the post-translational stage.

The ST/SC system was originally developed to produce synthetic peptides (33). It is based on spontaneous formation of an amide linkage between Lys and Asn, which leads to intramolecular isopeptide bond in the stable pilin protein in *Streptococcus pyogenes* (41, 42). After optimization, ST was determined to be a 13 aa peptide, which binds fast and stably with 138 aa SC under variant conditions (34, 35). Unlike antibody-antigen recognition, ST binds SC through an irreversible covalent linkage, which prompts its usage for protein modification and fluorescent imaging (43, 44). In this study, EsxA-ST was properly expressed and secreted to the mycobacterial surface where it was covalently modified with SC-GFP (**Fig. 4**). Covalent linkage of SC-GFP to EsxA-ST inhibited its MPA in liposome leakage assay (**Fig. 5**) and endogenous expression of SC-GFP inhibited Mm(EsxA-ST) intracellular survival in A549 cells (**Fig. 6**). Based on our experience, the rate of transient transfection and the rate of infection never reach 100% in the target cells. Thus, there were cells that were transfected but not infected and vice versa. In order to accurately measure the effect of SC-GFP expression on mycobacterial infection, we applied two independent approaches, fluorescence quantification and CFU, as described in **Fig. 6**. The fluorescence quantification is to calculate red/green ratio in the cells with both red (mCherry) and green fluorescence (SC-GFP or GFP). However, the CFU approach is not able to distinguish the heterogeneity of the cells. Because the cells that were infected but not transfected by SC-GFP were included into the CFU counting, the SC-GFP specific inhibitory effect was diluted, which may explain why at 24 hpi SC-GFP didn’t show significant inhibition on Mm(EsxA-ST) intracellular survival (**Fig. 6C**). While the overall inhibitory effect of SC-GFP on Mm(EsxA-ST) intracellular survival is significant when it is compared to GFP, the inhibition rate is only ~30%. The inhibition could be affected by the expression level and subcellular localization of SC-GFP (**Fig. S3**). Current studies suggest a model that mycobacteria are internalized into the phagosomes and EsxA mediates mycobacteria to escape from the phagosome and translocate to the cytosol for replicating and cell-to-cell spreading. Moreover, EsxA has been implicated to crosstalk with other cellular organelle and immune signal pathways (17, 45, 46), thus it is not clear at which steps and subcellular locations SC-GFP encounters and modifies EsxA-ST, which warrants further investigation. We also don’t exclude the possibility that EsxA only plays a partial role in phagosome rupture and cytosolic translocation, and other factors (e.g. Phthiocerol dimycocerosates) may also contribute to this process (17, 23, 47).

In summary, using DAS4+ system and ST/SC system we were able to knockdown EsxB and EsxA at the post-translational level and inhibit EsxA MPA at the post-secretional level. The results conclusively elucidate the direct role of EsxA MPA in phagosome rupture and cytosolic translocation. The two systems can be powerful tools in studies of host-pathogen interaction, gene function, protein-protein interaction and protein intracellular trafficking, etc.

## Materials and methods

### Bacterial strains, cell lines, plasmids and primers

The *Mycobacteria marinum* M strain, human lung epithelial WI-26 VA4 cell line and human monocyte THP-1 cell line were purchased from American Type Culture Collection (ATCC, Manassas, VA, USA) and preserved in our lab. Human lung epithelial A549 cell line was kindly provided by Dr. Jianying Zhang at The University of Texas at El Paso.

The pJSC407, pGOAL17 and pGMCKq1-10M1-sspBopt plasmids were kindly provided by Dr. Hugues Ouellet at The University of Texas at El Paso. The pJSC407 and pGOAL17 plasmids were used to produce the pJSC407-sacB suicide plasmid. The pGMCKq1-10M1-sspBopt was used for inducible expression of adaptor protein SspB for DAS4+ inducible knockout system. The pQE80L-SpyCatcher-ELP-GFP plasmid was purchased from Addgene (#69835, Watertown, MA, USA) for prokaryotic expression and purification of SpyCatcher-ELP-GFP protein.

We found that the purified SpyCatcher-ELP-GFP protein was toxic to human cells, which is presumably due to the cytotoxic effect of ELP linker between SC and GFP (48). Therefore, we removed ELP link to express SpyCatcher-GFP (SC-GFP) in *E. coli*. For eukaryotic expression of SC-GFP, we inserted SC into pcDNA3-EGFP vector. However, we found that transient transfection of pcDNA3-SC-GFP had a very low expression of SC-GFP in several mycobacteria susceptible cell lines, including WI-26, RAW264.7, THP-1 and A549. In order to enhance SC-GFP expression in eukaryotic cells, the coding sequence of SC was optimized for expression in human cell lines and synthesized by ThermoFisher Scientific (Waltham, MA, USA). The codon-optimized SC sequence was inserted into pcDNA3-EGFP (#13031, Watertown, MA, USA) for eukaryotic expression of SC-GFP. Even after optimization, SC-GFP was efficiently expressed only in A549 cell line.

The primers for site-directed mutagenesis, identification and RT-qPCR were all designed according to Mm’s genomic DNA sequence (GenBank Accession Number: CP000854.1) and synthesized by Sigma-Aldrich (St. Louis, MO, USA). The plasmids and primers used in this study are listed in **Table 1**.

**Table 1.**
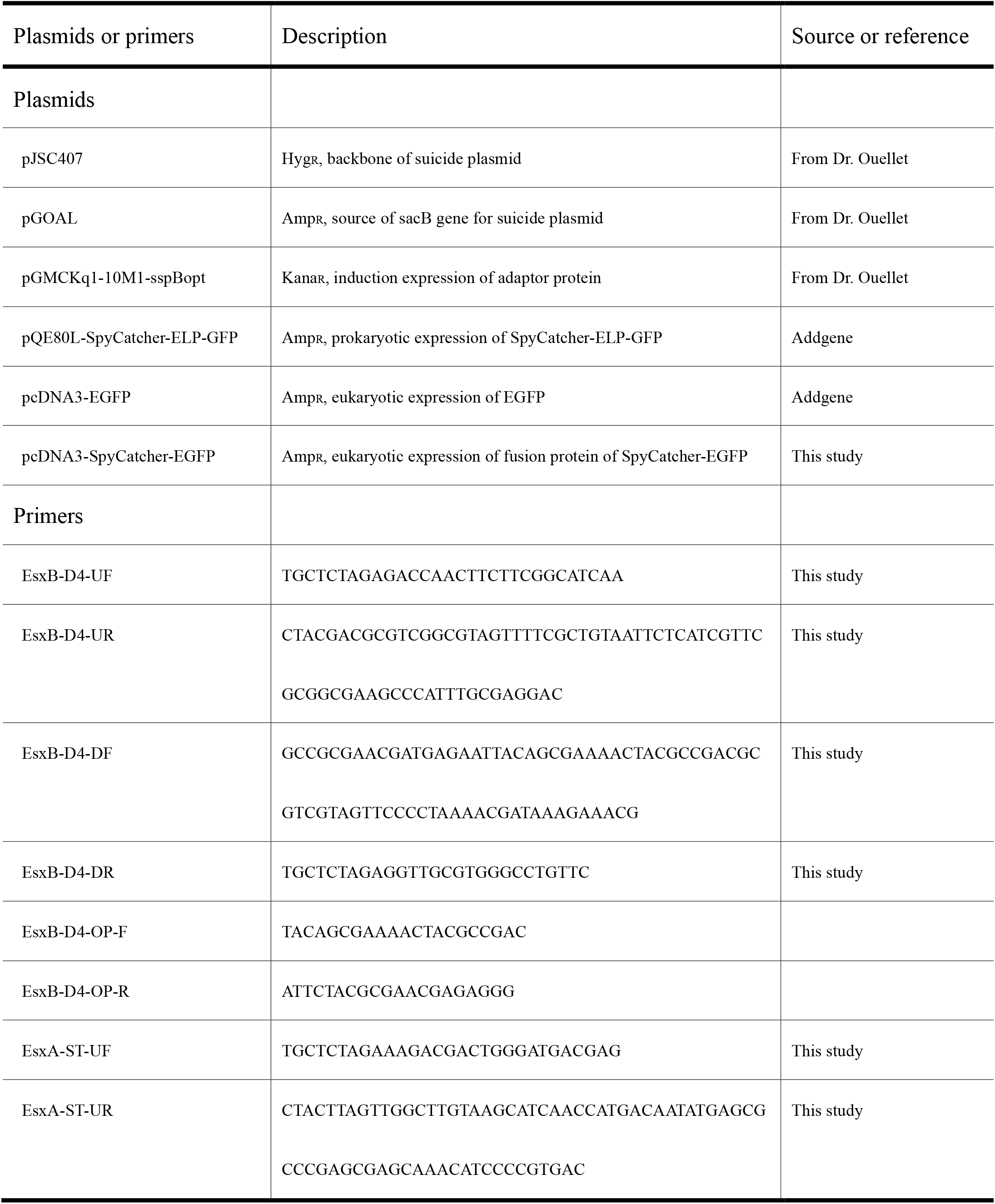
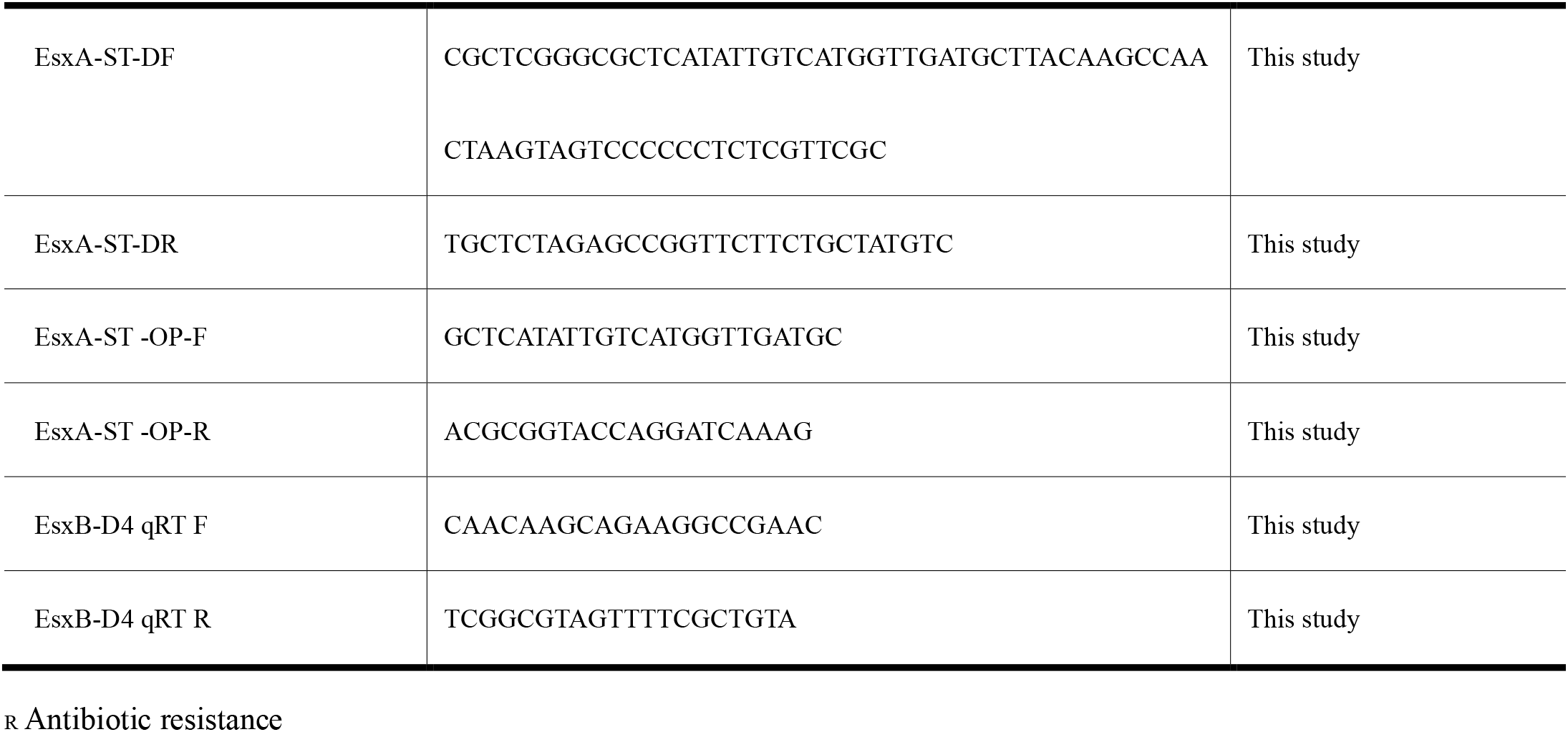
Plasmids and primers used in this study

To fuse ST or DAS4+ to the C-terminus of EsxA or EsxB, the homologous arms that flank the genes *esxA* or *esxB* were amplified from the genomic DNA of Mm, respectively. Then the amplified fragments were inserted into pJSC407-sacB. Similarly as previously described (21), the recombinant suicide plasmid was electroporated into competent Mm. After selection with 2% sucrose on the 7H10 plates (10% OADC), the colonies were picked, cultured and identified with primers that span the target genes (*esxA or esxB*) and the tags. The positive colonies were named as Mm(EsxB-DAS4+) and Mm(EsxA-ST), respectively. The plasmid pGMCKq1-10M1-sspBopt was electroporated into Mm(EsxB-DAS4+) to obtain Mm(EsxB-DAS4+)|pGMCKq1 for inducible knockdown of EsxB protein by ATC. All the Mm strains used in this study carry a mCherry-coding plasmid.

### Western blot and immunofluorescence assay

To detect the inducible knockdown of EsxB, Mm(EsxB-DAS4+)|pGMCKq1 was first cultured in 7H9 media (10% OADC, 0.05% Tween-80) until OD_600_ reached 0.8~1.0. Then, the culture was diluted again in 7H9 (10% OADC, 0.05% Tween-80) with 500 ng/mL of ATC at OD_600_=0.02. The culture pellet was then collected at various times of post induction. The expression of EsxB and EsxA in the total lysate was detected with Western blot by using anti-EsxB polyserum (NR-19361, BEI Resources, Manassas, VA, USA) and anti-EsxA polyserum (NR-13803, BEI Resources, Manassas, VA, USA).

Immunofluorescence assay was also used to test the inducible knockdown of EsxB-DAS4+. As described above, the Mm(EsxB-DAS4+)|pGMCKq1 pellet was collected at 48 h of post ATC induction. After the bacteria were fixed to the coverslip, the bacteria were incubated with 2% BSA for 30 mins at RT. After that, anti-EsxB polyserum and anti-EsxA polyserum were used as a primary antibody, and the FITC labeled secondary antibody was used to visualize bacteria-associated EsxB-DAS4+ and EsxA under confocal microscopy. To quantify the amount of bacteria-associated EsxB-DAS4+ and EsxA, the layers of FITC (green) and mCherry (red) were extracted from each image to calculate the overlap rate between the green and red signal. Briefly, the areas with green or red signals were first calculated. Then, the area of green that was colocalized with red was calculated. This area’s percentage in image’s total red area is the overlap rate, which was calculated according to equations in **Table 2**. Layer extraction and area calculation were achieved with Python 3.7.3 (49).

**Table 2.** Equations used for fluorescence image analysis

|  |  |
| --- | --- |
| Knockdown of EsxB-DAS4<sup>+</sup><sub>a</sub> | $\frac{areaG \cap areaR}{areaG}$ |
| SC-GFP inhibition on Mm(EsxA-ST) | $\frac{areaR \cap areaG}{areaG}$ |

a: areaG, area of green fluorescence signal; areaR, area of red fluorescence signal.

To detect the covalent bonding between SC-GFP and EsxA-ST, the total lysate of Mm(EsxA-ST) was incubated with purified SC-GFP at RT for 45 mins. The mixture was applied to SDS-PAGE and transferred to PVDF membrane, which was followed by Western blots using anti-GFP McAb (4B10, Cell Signaling Technology, Danvers, MA, USA) and anti-EsxA polyserum (NR-13803, BEI Resources, Manassas, VA, USA), respectively.

To label the bacteria-associated EsxA-ST with SC-GFP, the culture pellet of Mm(EsxA-ST) was collected and incubated with purified SC-GFP at RT for 45 mins. After several washes with PBS, the pellet was added into a 24-well plate with round coverslips. After a brief centrifugation, the bacteria were attached to the coverslips and then fixed with 4% paraformaldehyde. The fixed bacteria were observed with confocal microscopy (LSM 700, Zeiss, San Diego, CA, USA). The Mm(WT) was used as a control.

To test the effects of the endogenously expressed SC-GFP on Mm(EsxA-ST) intracellular survival, A549 cells were transiently transfected with pcDNA3-SC-EGFP or pcDNA3-EGFP as a control. At 24 h post transfection, the cells were infected with Mm(WT) or Mm(EsxA-ST) at MOI=2. At 24 hpi and 48 hpi, the cells in the randomly selected fields were imaged and the intracellular survival was calculated as the ratio of mCherry (red): EGFP (green). Same as above, the areas with green or red signals were first calculated. Then, the red areas that were colocalized with green were calculated. This area percentage in the total green area is the overlap rate we need, which was calculated according to equations in **Table 2**. Layer extraction and area calculation were achieved with Python 3.7.3 (49).

### Liposome leakage assay

As previously described (21), the liposome was produced from 1, 2-Dioleoyl-sn-glycero-3-phosphocholine (DOPC) lipid film. To encapsulate dye/quencher pairs, 8-aminonapthalene-1,3,6 trisulfonic acid (ANTS)/p-xylene-bis-pyridinium bromide (DPX) and DOPC film were rehydrated in 50 mM HEPES (pH 7.3) for six freeze-thaw cycles. Then, the mixture was filtered through 200 nM polycarbonate membrane (Avanti Polar Lipids, Alabaster, AL, USA) for 20 times. After that, the redundant salt in mixture was removed in a G25 desalting column (GE Healthcare Life Sciences, Pittsburgh, PA, USA) equilibrated with 50 mM HEPES solution (pH 7.3).

The dequenching of ANTS fluorescence was measured in an ISS K2 multiphase frequency and modulation fluorometer (ISS, Champaign, IL, USA) with excitation at 380 nm and emission at 520 nm. 100 μL of liposome and 100 μg of protein were mixed with the assay buffer (150mM NaCl, 20mM Tris, pH7.4) to a final volume of 1350 μL. After 30 s of incubation, 150 μL of 1 M NaAc solution (pH 4.0) was added into the mixture to activate acidic pH-dependent membrane insertion, and the fluorescence intensity was recorded continuously for the following 180 s. The mixture was continuously stirred throughout the assay. Prior to the assay, EsxA-ST was incubated with SC-GFP (molar ratio 1:0.5) for 2 hours at RT to allow formation of the covalent bond between EsxA-ST and SC-GFP.

### RNA extraction and RT-qPCR

The Mm(EsxB-DAS4)|pGMCK1q cells were treated with or without ATC and were collected from 5 mL liquid culture at 24 and 48 h post induction. Total RNA was extracted with TRI reagent (Ambion, Carlsbad, CA, USA). Total RNA was subjected to reverse transcription with High Capacity cDNA Reverse Transcription Kit (Thermo Fisher Scientific) at 25 °C for 10 min, 37 °C for 120 min, then 85 °C for 5 min for cDNA templates. SYBR_®_ Select Master Mix kit (Invitrogen) was used for RT-PCR, according to the manufacturer’s instruction: 1 μL cDNA, 1 μL forward or backward primer (10 μM), 7 μL nuclease-free water and 10 μL SYBR master mix were added. Reactions were on the Step One Real-Time PCR System (Applied Biosystems, USA) at 95 °C for 20 s, 40 cycles at 95 °C for 3 s, 60 °C for 30 s and a melting curve. Genes were tested in triplicate and the GAPDH gene was used as the internal control. Primers (EsxB-DAS4+ qRT F/R, Table 1) were designed to span the regions of the coding sequences of EsxB and the DAS4+ tag, based on National Center for Biotechnology Information (NCBI) Primer-BLAST (50). Relative transcription levels were calculated with the 2_−ΔΔCt_ method (51).

### Mycobacterial intracellular survival

To test mycobacterial intracellular survival in RAW264.7 cells, the cells were planted in 24-well plates at 5 ×10_5_ cells per well and cultured for 24 h. Then, mycobacterial single cell preparation was added into each well at MOI=1. After a brief centrifugation, the plates were incubated at 30 ℃, 5% CO_2_ for 30 min. After that, cells were washed with PBS to remove free bacteria. Dulbecco’s Modified Eagle Medium (DMEM) with 100 μg/mL Amikacin and 2% fetal bovine serum (FBS) was added to kill extracellular mycobacteria for 2 h. Then, the media was replaced with DEME containing 50 μg/mL Amikacin, 2% FBS and cultured at 30 ℃, 5% CO_2_. At 24, 48 and 72 hpi, the cells were lysed with 0.1% Triton X-100 and spread on the plates for colony forming unit (CFU) counting.

For mycobacterial intracellular survival in THP-1 cells, THP-1 cells were planted in 24-well plates at 1× 10_6_, then it was stimulated with 100 nM of 12-O-Tetradecanoylphorbol-13-acetate (PMA) overnight (52). After that, the differentiated THP-1 cells were rested for 24 h before they were infected with mycobacteria at MOI=1 for 4 h. To determine the effects of EsxB-DAS4+ knockdown on mycobacteria during infection, ATC was added into the experimental groups (5 μg/mL) to induce EsxB-DAS4+ degradation at the beginning of infection (0 hpi). The CFU of each group was determined at 24 and 48 hpi. For intracellular survival on WI-26, the cells were plated in 24-well plates at 2×10_5_ cells per well and cultured for 24 h. The Mm(EsxB-DAS4+)|pGMCKq1 cells were treated with ATC as described above. The pellet of both induced and uninduced groups were collected to produce single cell preparation. WI-26 cells were infected the same way as RAW264.7, the CFU of each group was determined at 48 h. To determine whether EsxB’s downregulation during intracellular infection would impact mycobacteria’s survival, WI-26 cells were infected as before. Then, ATC was added into media (5 μg/mL) at 0 hpi to induce EsxB-DAS4+ knockdown. CFU of each group was determined at 24 and 48 hpi.

To test mycobacterial intracellular survival in A549 cells, the cells were plated in 24-well plates at 1×10_5_ cells per well and cultured for 24 h. Same as above, the cells were transiently transfected with pcDNA3-SC-EGFP and pcDNA3-EGFP. The cells were then infected with Mm(WT) and Mm(EsxA-ST) at MOI=2. At 24 and 48 hpi, the cells were lysed with 0.1% Triton X-100 to determine the intracellular survival as CFU.

## Acknowledgement

The authors appreciate Hugues Ouellet for providing the pMSP12::mCherry plasmid, Jianying Zhang for providing A549 cell line and Xinzhuo Zhao for writing Python script of fluorescence images analysis. The study is supported by the grants from NIGMS (SC1GM095475 to J. Sun), National Center for Research Resources (5G12RR008124) and National Institute on Minority Health and Health Disparities (G12MD007592). The content is solely the responsibility of the authors and does not necessarily represent the official views of the National Institutes of Health. All authors have read the journal’s policy on disclosure of potential conflicts of interest, and the authors have no conflicts of interest to disclose. All authors have read the journal’s authorship agreement and the manuscript has been reviewed by and approved by all named authors.

## Supporting Information

**Fig.S1.**
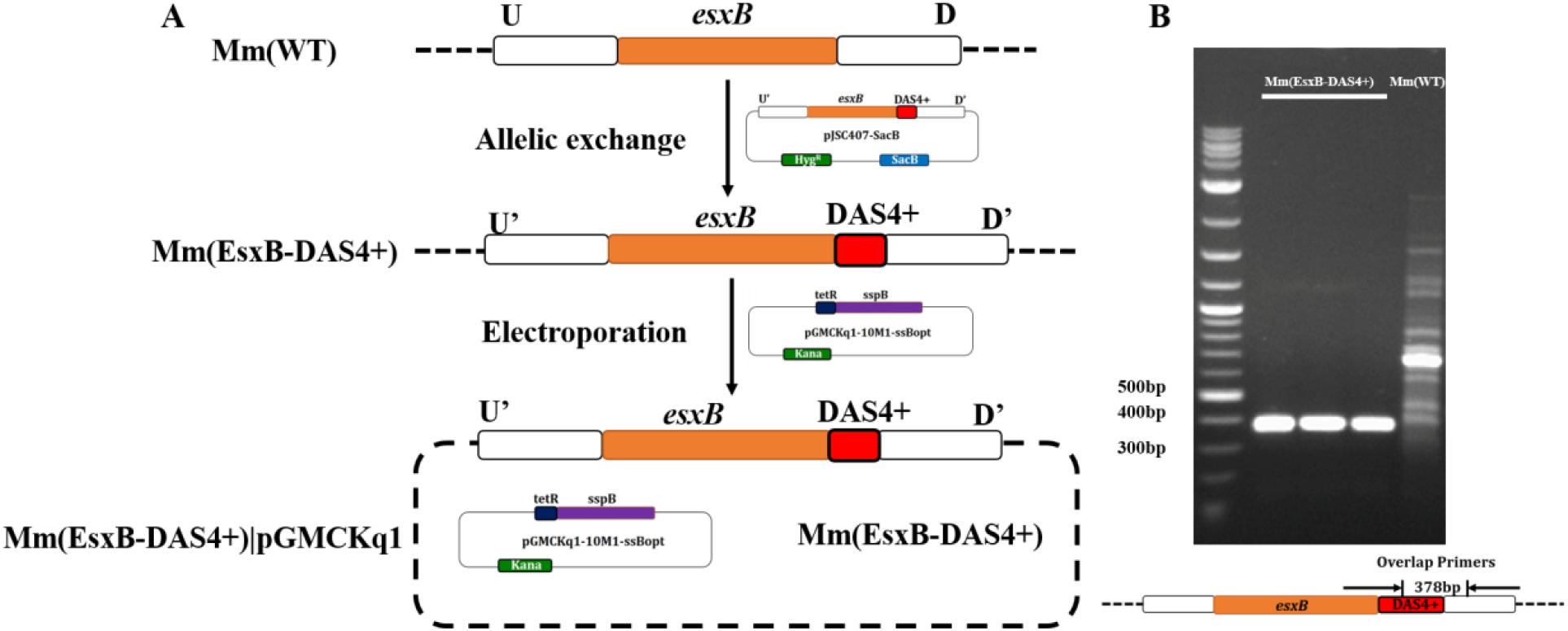
Construction and identification of the Mm(EsxB-DAS4+) strain. (**A**) The suicide plasmid pJSC407-sacB was used to insert DAS4+ to the C-terminus of EsxB. Then the pGMCKq1-10M1-ssBopt plasmid that encodes adaptor protein was electroporated into Mm(EsxB-DAS4+). (**B**)The genomic DNA of Mm(EsxB-DAS4+) was extracted and applied to PCR to confirm the insertion of DAS4+. The primers that overlap the insert sequence were used for PCR and produced a specific DNA fragment with an expected length ~ 400 bp, which is absent in the genomic DNA extracted from Mm(WT).

**Fig.S2.**
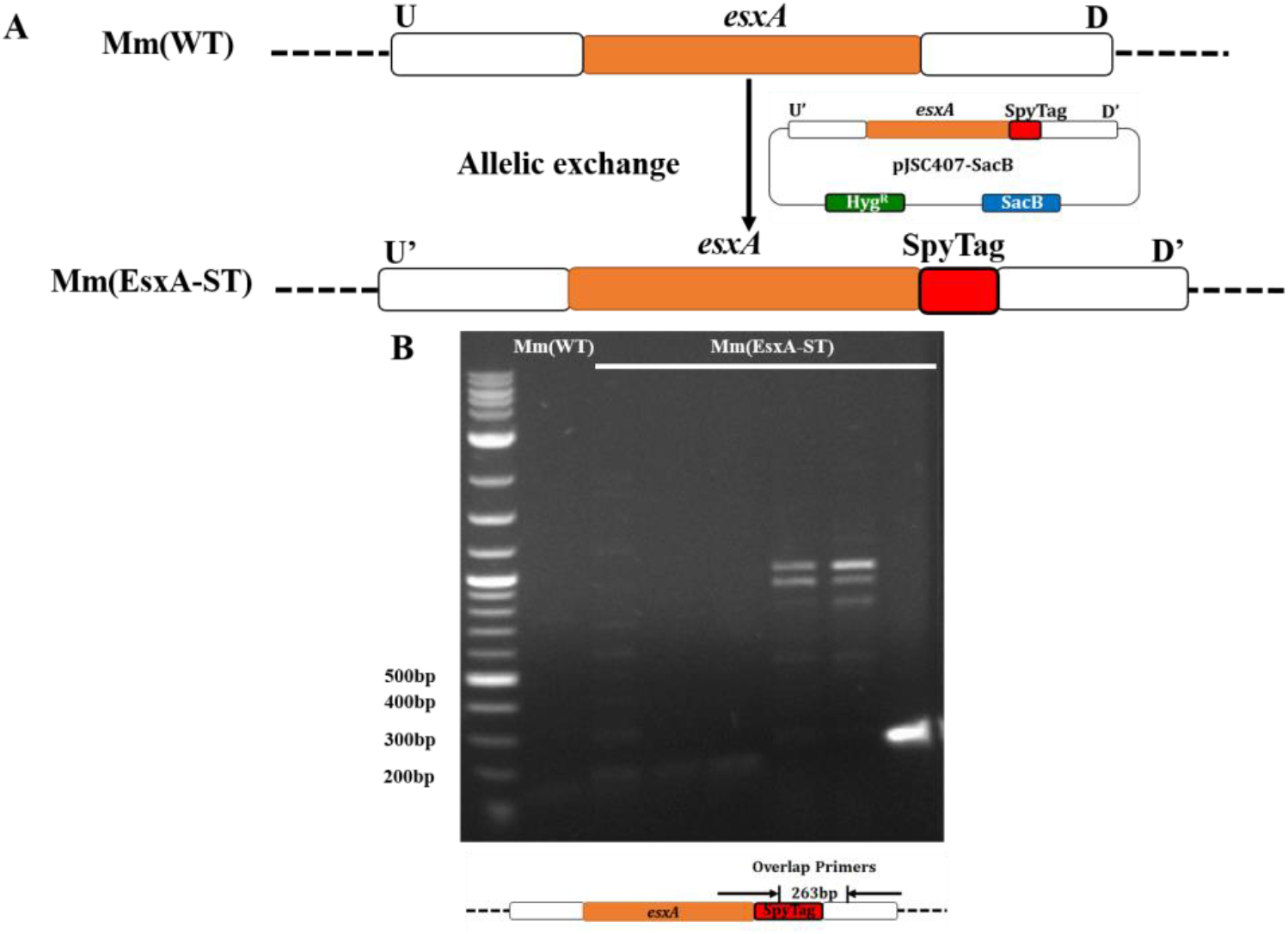
Construction and identification of the Mm(EsxA-ST) strain. (**A**) The suicide plasmid pJSC407-sacB was used to attach ST to the C-terminus of EsxA. (**B**) The genomic DNA from several clones of the Mm(EsxA-ST) candidates was extracted and applied to PCR with the primers that overlap the insert sequence to confirm the insertion of ST. One of the clones produced a specific DNA fragment with an expected length ~ 300 bp, which is absent in the genomic DNA extracted from Mm(WT).

**Fig.S3.**
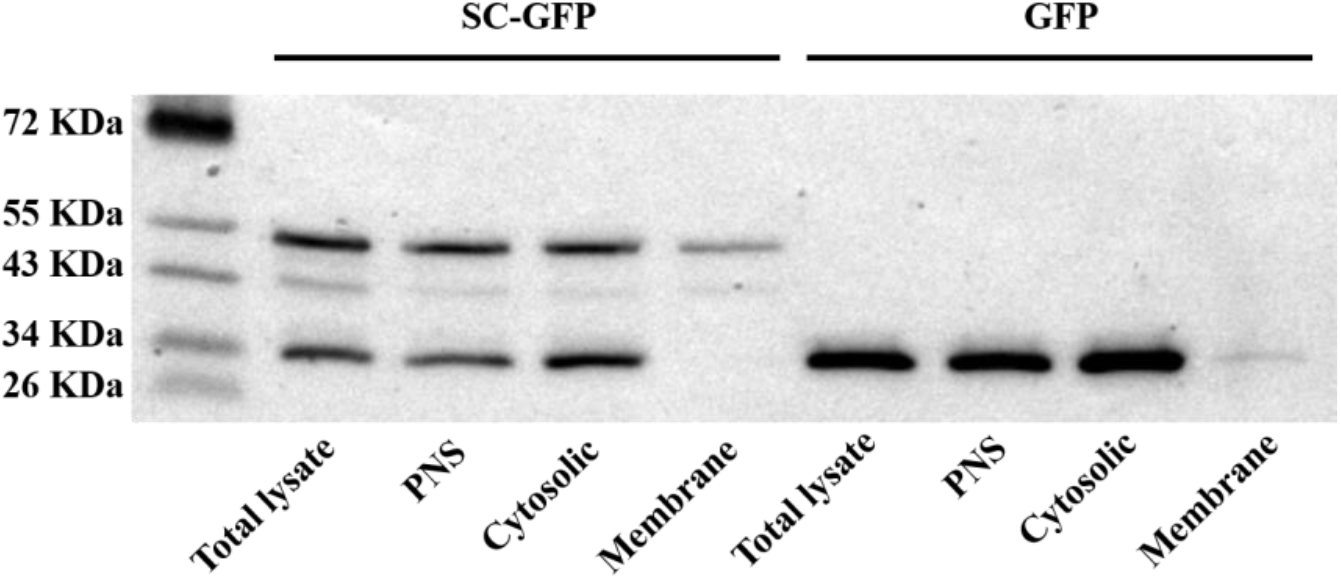
Subcellular fractionation of SC-GFP in A549 cells. The A549 cells were transfected with pcDNA3-SC-EGFP or pcDNA3-EGFP for 24 hours and then the cells were harvested, lysed and fractionated into post-nuclear supernatant (PNS), cytosolic fraction and membrane fraction. The samples were applied to SDS-PAGE, followed by Western blots using the antibody against GFP.

